# Identification of interacting proteins of maize mosaic virus glycoprotein in its vector, *Peregrinus maidis*

**DOI:** 10.1101/2022.02.01.478665

**Authors:** Karen Barandoc-Alviar, Dorith Rotenberg, Kathleen M. Martin, Anna E. Whitfield

## Abstract

Rhabdovirus glycoproteins (G) serve multifunctional roles in virus entry, assembly, and exit from animal cells. We hypothesize that maize mosaic virus (MMV) G is required for invasion, infection, and spread in *Peregrinus maidis*, the planthopper vector. Using a membrane-based yeast two-hybrid assay, we identified 125 *P. maidis* proteins that physically interacted with MMV G, of which 68% matched proteins with known functions in endocytosis, vesicle-mediated transport, protein synthesis and turnover, nuclear import/export, metabolism and host defense. Physical interaction networks among conserved proteins indicated a possible cellular coordination of processes associated with MMV G translation, protein folding and trafficking. Non-annotated proteins contained predicted functional sites, including a diverse array of ligand binding sites. Cyclophilin A and apolipophorin III co-immunoprecipitated with MMV G, and each showed different patterns of co-localization with G in insect cells. This study describes the first protein interactome for a rhabdovirus spike protein and insect vector.

## Introduction

Virus infection of host cells is a coordinated process mediated by interactions between viral and host proteins. Viral spike proteins are the first proteins to encounter host cell surfaces. Such is the case for members of the *Rhabdoviridae*, a large virus family composed of animal- and plant-infecting negative-sense RNA viruses encapsidated in complex virions. Rhabdoviruses encode a single spike protein (G) that projects from the surface of the particle and plays a role in viral attachment and fusion (reviewed in Belot et al., 2019). Much of what is understood about rhabdovirus G protein - host interactions comes from molecular investigations of animal-infecting members of the family, notably *Rabies virus* (RABV), *Vesicular stomatitis virus* (VSV) and *Viral hemorrhagic septicemia* (VHSV) (Bearzotti et al., 1999; Belot et al., 2019). Some of the host cell receptors identified for rhabdoviruses include low density lipoproteins receptor (LDL-R) family members (Finkelshtein et al., 2013), nicotinic acetylcholine receptor (Gastka M. 1996; Lentz 1982), integrin β1 (Shuai et al., 2020), and fibronectin (Bearzotti et al., 1999). Reports implicate other host proteins and membrane structural components as putative receptors for RABV, VSV and VHSV (Carneiro et al., 2002; Schlegel et al., 1983; Thoulouze et al., 1998). Given the remarkable diversity of rhabdoviruses and their respective hosts and tissue tropisms, the characterized receptors are similarly varied and may be unique for a given species of virus within the family. In contrast, the entry process is conserved and uses the clathrin-mediated endocytic pathway followed by low-pH-induced membrane fusion (Belot et al., 2019). G undergoes structural rearrangement from its pre- to post-fusion conformation during this fusion process (Belot et al., 2019; Da Poian et al., 2005). The G protein is also essential in virus particle assembly as it is a major structural component, and it may have additional host-protein interaction partners involved in nuclear import and export for the nuclear-replicating plant rhabdoviruses. While animal-infecting rhabdoviruses have served as models for G protein-mediated host cell entry for uncharacterized rhabdoviruses (Roche et al., 2006, 2007; Yang et al., 2020), the interactions between the G protein of plant- and insect-infecting rhabdoviruses with insect-vector proteins are virtually unknown.

*Maize mosaic alphanucleorhabdovirus* is a plant-pathogenic rhabdovirus that is transmitted in a persistent propagative manner by the corn planthopper, *Peregrinus maidis*. The virus and the vector are pests in tropical and subtropical climates and cause yield losses in important staple food crops like maize and sorghum (Dietzgen et al., 2020; Singh and Nadoor, 2008). The vector sustains a lifelong virus infection with little to no pathology, resulting in efficient and long-term transmission (Barandoc-Alviar et al., 2016; Higashi and Bressan et al., 2013). Maize mosaic virus (MMV) can be acquired efficiently by all five nymphal stages and both sexes of adult *Peregrinus maidis*, and older nymphs and adults are capable of transmission to plants after a long latent period of three to five weeks (Ammar and Hogenhout, 2008). When acquired as nymphs, virus titers steadily increase over development and peak at the adult stage (Barandoc-Alviar et al., 2016).

The MMV/*P. maidis* is a model system for understanding virus-vector interactions for plant rhabdoviruses (Redinbaugh and Hogenhout, 2005; Whitfield et al., 2018; Yao et al., 2013, 2019). The virus exhibits broad tissue tropism in the vector, replicating in the midgut, trachea, salivary glands and neural tissues (Ammar and Hogenhout, 2008; Yao et al., 2019). The infection pathway begins at the midgut epithelium, disseminating to other cell types and organs. Detection of MMV in hemocytes and nerve cells indicates hemolymph or neural dissemination routes in the planthopper vector (Ammar and Hogenhout, 2008; Yao et al., 2019). Widespread infection of *P. maidis* adult bodies culminated in transcriptome-wide, differential responses to MMV, including processes associated with protein and nucleotide binding, transmembrane transport and innate immunity (Martin et al., 2017). At the cellular level, MMV replicates in the nucleus of the insect vector cells and MMV protein localization patterns have been mapped in insect cells and demonstrated that G primarily localizes to the nuclear membrane with some protein detected in the cytoplasm that may be associated with the ER (Ammar et al., 2009; Martin and Whitfield, 2018).

Proteomic methods have yielded significant advancements towards understanding the mechanisms of virus-vector interactions (Rajarapu et al., 2021; Wilson et al., 2020). Omics technologies have enabled new biological insights into mechanisms of virus replication, virus entry, antiviral host responses and viral subversion of host defenses. Throughout the course of infection, host cellular mechanisms rely heavily on the formation of temporally and spatially regulated virus–vector protein–protein interactions. Here, we used the membrane-based yeast two-hybrid (MbY2H) system to investigate interactions of the MMV surface glycoprotein G, an integral membrane protein. MbY2H is based on split-ubiquitin protein complementation assay and detects protein interactions directly at the membrane, thereby allowing the use of full-length G to screen a *P. maidis* cDNA library to identify host protein interaction partners. We confirmed, sequenced and reduced redundancy in a set of putative MMV G interactors with diverse annotations consistent with mechanisms associated with virus entry, intracellular transport, and host defense, and a subset of non-annotated interactors with predicted functional sites. In insect cells, we validated the interactions between G and proteins with established roles in virus-host interactions (cyclophilin A, CypA) and hemolymph lipid transport (apolipophorin III, ApoLpIII). Further characterization of the planthopper proteins will lead to new understanding of rhabdovirus-host interactions.

## Materials and Methods

### Split ubiquitin membrane-based yeast two hybrid assay (MbY2H)

We used the Dualsystems (Zurich, Switzerland) DualMembrane Starter Kit for in vivo detection of membrane-protein interactions in the budding yeast *Saccharomyces cerevisiae*. For the screen, the viral glycoprotein was used as bait and a *P. maidis* cDNA library as prey. Laboratory colonies of *P. maidis* and MMV-infected plants were maintained as previously described (Barandoc-Alviar et al., 2016). All procedures leading to library screening were done following the manufacturer’s procedures for bait construction, verifying expression of bait in yeast, validating true interactions by re-transformation with the bait in the yeast reporter strain, functional assay of the bait through interaction with a known protein, and self-activation test of the bait. We performed a pilot MbY2H screen to optimize the stringency of selection in MbY2H library screening.

### Construction and optimization of the bait, MMV-G

MMV G was cloned into the yeast bait plasmid, pBTR3-SUC, using primers described in **Supplementary Table S1**. Prior to use in the library screen, correct expression of the bait fusion protein, pSUC-MMV G in yeast was determined using a functional assay where the bait was transformed with a control prey plasmid pOst1-NubI. To verify that the bait did not self-activate, we tested different stringency of 3-AT in the selection plates. Based on the growth or no growth of yeast on the selection plate, Synthetic defined medium lacking leucine, tryptophan, histidine and adenine (SD-LWHA) amended with 1 mM 3-AT was selected for the library screen.

### cDNA preparation for directional cloning into the prey vector

The cDNA library was made from a pool of RNA samples from different developmental stages of *P. maidis* from third nymph (N3) to adult (1-3 days after N5 molt to adult) and insects were reared on MMV-infected plants (Barandoc-Alviar et al., 2016). The quality of RNA from each stage was analyzed using Agilent Bioanalyzer’s RNA 600 Nano assay class (Agilent Tech., Sta. Clara, CA) before pooling equal amounts to be used for cDNA synthesis. Full-length enriched double-stranded (ds) cDNA was synthesized using EasyClone cDNA library construction kit (Dualsystems Biotech) following manufacturer’s protocol. Briefly, 2 ug of pooled total RNA was used for first strand cDNA synthesis. The synthesized first strand cDNA was used to amplify ds cDNA by long distance PCR (LD-PCR). After amplification, the cDNA was purified using QIAquick PCR purification kit (Qiagen, Valencia, CA) and digested with *Sfi*I. Following the company’s protocol, the sample was size fractionated using CHROMA SPIN 1000 columns to select for DNA sizes greater than 1 kb (Clontech, Mountain View, CA). After fractionation, the concentration of cDNA was determined using Nanodrop ND 1000 spectrophotometer (Thermo Fisher Scientific, Lenexa, KS) and its size and quality were checked using Agilent Bioanalyzer’s DNA 7500 assay class. The full-length enriched cDNA library after fractionation was checked for quality and size distribution before it was directionally cloned into the prey vector, pPR3-N (Supplementary Figure 2). pPR3-N was digested with *Sfi*I and after linearization, it was purified using QIAquick gel purification kit (Qiagen, Valencia, CA) and used for ligation with the digested cDNA. The ligation reaction was composed of 500 ng of library vector and 200 ng of cDNA. Transformation with Electromax DH10B cells was done using 1 ul of the ligation reaction mixture. Afterwards, the efficiency of the transformation reaction and the titer of the library were determined.The quality assessment of the library revealed a library titer of 4 × 106 CFU/ml and the average insert size was 1.5 kb. Bait self-activation assays were conducted, and we determined that 1 mM 3-AT was sufficient to prevent self-activation of MMV G.

### Amplification of the cDNA library

The ligation mix was transformed into a high efficiency electrocompetent cells, DH10B Electromax (Invitrogen, Carlsbad, CA). About 120 large (150 × 15 mm) LB plates supplemented with 100 ug/mL ampicillin were used for amplification. The cells were harvested and re-grown for 3 h at 30°C with vigorous shaking. From the 3 L culture, 30 ml of culture was removed, and the cells were collected and resuspended in 10 ml LB supplemented with 25% glycerol. The cells were aliquoted in 1 ml into screw cap cryotubes and quickly frozen in liquid nitrogen and stored in -80 °C and this served as the primary library stock. The remaining bacterial cell culture was prepared for plasmid DNA isolation using Qiagen Plasmid Plus midi kit (Valencia, CA).

### Screening of a P. maidis cDNA Library using the split-ubiquitin MbY2H assay

We used the DUALmembrane kit (Dualsystems Biotech) to screen the *P. maidis* cDNA library. The genotype of host strains used are listed in **Supplementary Table S2**. The bait fusion vector pSUC-MMV G in NMY51 was transformed with the cDNA library using the lithium acetate method with ssDNA. The mixture was spread on the selection plate SD-LWHA + 1mM 3-AT and SD-LW and incubated for 4 days at 30 °C. The colonies were picked and re-streaked in the same selective medium for β –galactosidase enzyme activity assay. In addition to the growth reporters *HIS3* and *ADE2*, the yeast strain NMY51 has the color reporter *lacZ*. Picked colonies from the library screen were analyzed for β-galactosidase activity using the manufacturer’s protocols. Plasmid DNA was isolated from all the colonies and they were re-transformed into bacteria. Plasmid was isolated and digested with *Sfi*I restriction enzyme to determine the size of the prey. To verify if they were true interactors in the MbY2H assay, the plasmid DNA from bacteria was re-transformed with the bait in yeast in the same selection plate. The samples which grew in the selection plate were subjected to β-galactosidase activity assay to eliminate false positives. Colonies positive for this assay (n= 432) were picked for yeast plasmid isolation, re-transformed into *E. coli*, and sequenced using Sanger technology (**Table 1**). Iterative analysis of the sequences as they were generated revealed that sequencing of a subset of the validated clones (432 of 616) was approaching saturation in the diversity of sequences represented in the screen.

**Table 1.** Sequential outputs of the membrane-based yeast II hybrid screen (MbY2H) for protein-protein interactions between maize mosaic virus glycoprotein (MMV-G, bait) and a *Peregrinus maidis* cDNA library (prey).

| Procedural Step | Counts |
| --- | --- |
| Colonies picked | 1, 264 |
| <sup>a</sup>Putative interactors | 1, 091 |
| <sup>b</sup>Confirmed interactors | 616 |
| Samples sequenced (cDNA) | 432 |
| <sup>c</sup>Non-redundant cDNA sequences | 125 |

<sup>a</sup>colonies that tested positive for β-galactosidase activity; <sup>b</sup>re-transformed putative interactors that interacted with the bait (MMV-G) for a second round of MbY2H screen to remove false positives; <sup>c</sup>final set of interactors (cDNA) after filtering out redundant sequences with CD-HIT (v. 4.6.8) through the CYVERSE cloud computing resource (www.cyverse.org); criteria for non-redundancy = % nucleotide identity less than 93% between any two sequences

### Bioinformatic analysis of MMV-G interactors

To reduce redundancy among the 432 transcript sequences identified from the Y2H screen, the nucleotide sequences were subjected to a CD-HIT analysis (v. CD-HIT-est_4.6.8) through the CYVERSE cloud computing resource (www.cyverse.org) to cluster highly similar transcript sequences (global sequence %ID cutoff = 93%) and to identify the longest representative sequence from each cluster. Blastx and Gene Ontology (GO) analyses were performed on the reduced sequence (non-redundant) dataset on January 28, 2021 with OmicsBox 1.4.11 using default parameters. The distribution of GO-annotated sequences into their most specific GO terms was determined by performing a multilevel analyses in OmicsBox. In addition, Blastn analyses were performed locally on the non-redundant Y2H sequences (nucleotide queries) using two previously published transcriptome datasets for *P. maidis*: a de novo assembled contigs of RNAseq reads for adult intact bodies (68,003 contigs) (Martin et al., 2017) and expressed sequence tags (20,678 ESTs, Sanger sequencing) for nymphal (N4) guts (Whitfield et al., 2011). Both blastn comparisons were performed in OmicsBox using default settings with an E-value cut-off of 10^−10^.

As a means toward deeper understanding of the roles of G-interactors, a STRING analysis (Szklarczyk et al., 2021; https://string-db.org/) was performed to predict both physical and functional interactions between the non-redundant dataset. Open reading frames (ORFs, amino acids) were generated by NCBI ORF finder to identify the longest, complete ORF (start and stop codons) (https://www.ncbi.nlm.nih.gov/orffinder/) for each *P. maidis* G-interactor. The ORF sequences (100 out of 125) were mapped against a *Drosophila melanogaster* protein database, then interaction networks were generated with a minimum interaction score of 0.4 (medium confidence) to produce a physical interaction network and a network that included both functional and physical interactions between G-interactors.

To aid in annotation of G-interactors with no significant match to known proteins or those that matched hypothetical/unknown/uncharacterized proteins in other metazoans, Eukaryotic Linear Motif (ELM) was performed, a computational tool that predicts short linear motifs (SLiMs) that infer functionality (sites) in eukaryotic proteins (Dinkel et al., 2016; Kumar et al., 2020; http://elm.eu.org/index.html). Sequences that occurred in CD-HIT clusters (i.e. groups of highly similar sequences, at least 93% identical at the nucleotide level) were selected to limit the analysis to sequences with higher confidence (meaning, more than one clone for a given sequence identified from the Y2H screen). This reduced the ELM dataset to 17 non-annotated sequences. Predicted ORFs were run through ELM with the following analysis parameters: *Peregrinus* maidis as the specified organism, motif probability cut-off = 0.001 (≤ 10^−4^ = E), and motifs with low conservation were not reported. Functional class site, ELM class, ELM ID no. and accession, and ELM role were recorded for each of the sequences.

### GFP-, RFP-, and Myc- tagged plasmid constructs for co-immunoprecipitation and localization experiments

The coding sequence of MMV-G, CypA, and ApoLpIII were cloned into pENTR (Invitrogen, Carlsbad, CA, USA) by inserting a Kozak ATG initiation sequence to the forward primer to allow proper translation initiation. The reverse primer was designed in such a way that the PCR product was in frame with a C-terminal tag by removing the native stop codon in the gene of interest. Following cloning and sequencing, expression of GFP-, RFP-, and Myc-tagged genes in S2 cells were achieved by recombination of the pENTR constructs of the candidate genes (ORF) into the destination vectors having an HSP70 promoter using LR clonase II (Invitrogen, Carlsbad, CA, USA). The destination vectors were: pHWG (GFP tag at the C terminal of the inserted gene), pHRW (RFP at the N terminal end of the inserted gene and pHWM (Myc tag at the C terminal end of the inserted gene (Drosophila gateway collection, DGRC, Bloomington, Indiana). For colocalization experiments in Sf9 cells, the MMV-G, CypA and ApoLpIII entry clones were recombined into pIB vectors. pIB was modified to have GFP and RFP after the DEST cassette like previously reported with the exception of an AscI site on the 5’ end of the fluorophore (Maroniche et al., 2011). All expression constructs were sequence-verified before using in transfection experiments. The pENTR constructs were recombined into the pIB-v5-his-dest GFP and RFP vectors using LR clonase II.

### Co-immunoprecipitation assay

S2 cells were cultured in Schneider’s insect medium (Gibco) supplemented with 10% FBS (Gibco). MMV G-FLAG and Pm constructs were separately transfected into S2 cells using Cellfectin II transfection reagent (Life Technologies). The amount of each construct transfected into the cells was equivalent to ensure comparable expression levels of the proteins. Cells were harvested three days post transfection, mixed together with rotation for 1 hr at RT. Equivalent amounts of mock-transfected cells were used as control and the final mixtures contained the same amount of cells under all co-immunoprecipitation conditions. After incubation, the cells were lysed in Pierce IP Lysis buffer (Thermo Scientific, Waltham, MA) supplemented with protease inhibitor cocktail, PIC (Roche, Indianapolis, IN), phenylmethanesulfonyl fluoride, PMSF (Sigma-Aldrich, St. Louis, MO) and 0.5% Nonidet NP-40 (Amrresco, Solon, OH). Then, cell lysates were incubated with anti-FLAG affinity gel (Biotool, Houston, TX) overnight at 4 °C with rotation. After incubation, the samples were washed extensively with lysis buffer and the proteins were eluted by boiling 5 ul of the resin with 5x SDS loading buffer and the proteins were detected by western blot analysis using rabbit antibody to FLAG (1:2,000, BioRad) or mouse antibody to Myc (1:2000, Sta. Cruz) and Clean-Blot IP detection reagent (HRP) (1:5000, Thermo Scientific).

### Immunoblot Analysis of Proteins

Protein expression in S2 cells transfected with plasmid DNA was assessed by Western blot analysis three days post transfection. A total of 10 ml culture was lysed using Pierce IP Lysis buffer (Thermo Scientific, Waltham, MA) supplemented with protease inhibitor cocktail, PIC (Roche, Indianapolis, IN), phenylmethanesulfonyl fluoride, PMSF (Sigma-Aldrich, St. Louis, MO) and 0.5% Nonidet NP-40 (Amrresco, Solon, OH). Twenty-five ul of lysate was mixed in Laemmli sample buffer (60 mM Tris–Cl pH = 6.8, 2% SDS, 10% glycerol, 5% β-mercaptoethanol, 0.01% bromophenol blue). Proteins were separated using a gradient SDS PAGE gel (8-12%) and a Mini-Protean 3 Cell (Bio-Rad) at 85 V. Proteins were electroblotted as described by Badillo-Vargas et al. (Badillo-Vargas et al., 2012). The membranes were scanned using LI-COR Odyssey Imaging System (LI-COR Biotechnology, Lincoln, NE, USA) for detection of protein.

### MMV G localization in insect cells using organelle probes

To examine MMV G may be localization in insect cells, we used different organelle probes to locate MMV G expression after transfection in S2 cells. We used ER-Tracker Blue-White DPX and Lyso Tracker® (Life Technologies, Carlsbad, CA, USA) which are selective for endoplasmic reticulum and lysosomes, respectively. S2 cells were fixed on coverslips using 10% formaldehyde in PBS buffer (pH 7.2).After removing cell culture media, the cells were covered to a depth of 2– 3 mm with 10% formaldehyde diluted in warm PBS. The cells were fixed for 15 min. After fixing with formaldehyde, 1 ml of fresh PBS was added and they were transferred into a 12-well culture plate. The cells were allowed to sit for about 15 min. About ∼750 ul of the media were removed and 20 ul of the ER tracker dye was added to the cells dropwise. ER-Tracker Blue-White DPX staining solution was prepared by diluting a 1 mM stock solution to the lowest recommended final working concentration of 100 nM to minimize potential labeling artifacts. After incubation of 15 min, 5 ul of cells were put on the slide and covered with glass. The cells were viewed using a confocal microscope at excitation or emission (ex/em): 374/430-640. For staining lysosomes, LysoTracker Red was used at the suggested working concentration of 50 nM. This was prepared in S2 medium without FBS and stored in a dark-colored 0.5 ml microfuge tube. This dye was always prepared fresh RT (to 37 °C). After fixing and staining cells they were viewed under a confocal microscope at ex/em: 577/590.

### Transfection, Localization and Imaging of Planthopper Proteins

For sub-cellular localization studies of CypA and ApoLpIII in Sf9 cells, Lipofectamine LTX (Thermo Fisher Scientific, Waltham, MA, USA) was used for transfection of plasmid DNA in 90% confluent Sf9 cells (Vaughn et al., 1977) following the manufacturer’s recommendations. Cells were plated in a 12-well plate and had a concentration of 0.5× 10^5^ cells/ml for transfection. Each well was transfected with 1ug of plasmid DNA and 6 ul of Lipofectamine LTX and 6uL PLUS reagent in Grace’s media. The plate was incubated at RT overnight and the medium was replaced with fresh Sf9III medium. For co-transfection, equal amounts of plasmid DNA of GFP-tagged MMV G and RFP-tagged *Pm* genes, CypA and ApoLpIII were used. Two days after transfection, cells were stained with 105ng/uL of 4’,6-Diamidino-2-Phenylindole, Dihydrochloride (DAPI) in PBS for twenty minutes and then washed three times with PBS. For each construct, there were a minimum of two wells treated and visualized and the experiment was replicated three times to confirm localization patterns.

Transfected and stained cells were viewed using Zeiss LSM 880 inverted confocal laser scanning microscope (Zeiss, Oberkochen, Germany) using a C-Apochromat 40x/1.1 W Korr M27 objective and Zen black software v. 10.0.0 with a pixel average of 4 and 16 bit depth and accompanied by Zen lite software, blue version 3.0 to export images into Tiff format.

## Results

### Diverse coverage of putative G-interactors were identified by the MbY2H screen

We used the split-ubiquitin membrane-based yeast two hybrid system to identify potential planthopper proteins which interacted with MMV G. With the intention to recover G-interacting *P. maidis* protein from transmission-competent insects, we generated a cDNA library collectively capturing sequences from later nymphal stages (N3 - N5) and adults (females and males). For the MbY2H interaction screen, 1,264 colonies were picked, of which 1,091 were determined to be putative interactors of MMV G after testing positive for β-galactosidase activity (**Table 1**). After a second round of the interaction screen with the β-galactosidase activity-positive clones, we confirmed 616 putative interactors, and from this collection, we sequenced 432 of the clones. Annotation of the sequences revealed that many interactors were identified multiple times indicating that the 432 clones reflect the diversity of interactors that are possible in this MbY2H screen.

### Annotation of G-protein interactors predict unique proteins and enrichment in membrane-associated proteins

The CD-HIT analysis of the 432 transcript sequences resulted in a dataset of 125 non-redundant sequences with a median length of 1,014 nucleotides (**Supplementary Fasta sequence file**; **Supplementary Table S3**). Blastx analysis revealed that 32% of these 125 putative G-interactors had no significant match to proteins (non-annotated), possible indication of proteins unique to *P. maidis*, and 16.8% significantly matched putative proteins in other species that have yet to be functionally annotated (i.e., uncharacterized, or unknown or hypothetical). Of the 85 sequences with significant blastx protein matches, 80% were assigned GO terms that provisionally indicated their cellular roles (biological process and molecular function) and localization (**Supplementary Table S3**). While there was no apparent sequence enrichment of any particular GO term associated with a biological process or molecular function, GO terms with the highest number of sequences were ‘ATP synthase proton coupled transport’ (17%, **Supplementary Figure S1A**) and ‘ATP binding’ (14%, **Supplementary Figure S1B**). Most notably, however, 37% of the GO-annotated sequences in the multilevel analysis were enriched in the GO term ‘integral component of membrane’ (**Supplementary Figure S1C**). Likewise, manual examination of the GO terms assigned to *P. maidis* sequences that matched uncharacterized/unknown/hypothetical proteins in other species revealed that 76% of these (16/21) were GO-classified only by subcellular localization, all of which were predicted to be localized to membranes (**Supplementary Table S3**). As such, it appears that the set of *P. maidis* proteins that interact with MMV G are biased towards proteins localized to membranes.

### Transcripts-encoding putative G-interacting proteins are expressed in P. maidis bodies and guts

Whole body and gut transcriptome data previously published for *P. maidis* (Martin et al., 2017; Whitfield et al., 2011) were mined to confirm the presence of the transcripts that encode the 125 non-redundant, MMV G-interacting proteins, including those that were non-annotated (no protein match) or uncharacterized/unknown/hypothetical in other species. Approximately 85% of the 125 nucleotide sequences significantly matched *P. maidis* adult whole body transcripts (mean % similarity = 95.8%, mean *E* = 1.52 × 10^−25^), and ∼ 63% significantly matched nymph gut transcripts (mean % similarity = 92.8%, mean E = 92.86 × 10^−14^) (**Supplementary Table S4**). Of the 40 non-annotated (no match, putatively *P. maidis*-specific) G-interactors, 68% and 45% were found in the whole body and gut transcriptomes, respectively, confirming the validity of these sequences in the current project. Likewise, of the 17 uncharacterized/unknown/hypothetical proteins reported in other species, 76% and 62% of these sequences were expressed in whole bodies and gut transcriptomes of this planthopper, respectively. Overall, regardless of sequence database size, the analyses confirmed that the majority of the putative G-interactor sequences were accounted for in two independent studies.

### In silico analysis of protein-protein interactions among the G-interactors implicate protein translation, folding and trafficking

To gain insight into the possible cross-interactions among the 125 G-interactors of *P. maidis*, protein-protein interaction networks were constructed on evolutionarily conserved proteins. Of the 100 predicted ORF sequences, only 29 matched the proteome sequences of *D. melanogaster*, the closest species with a functionally-annotated proteome. Of these proteins, there are reports of experimental evidence that 14 interact physically with one or more members of this set of 14 in *Drosophila* (**Figure 1A**), and an additional three proteins have been reported to exhibit indirect, functional interactions (i.e., evidence of co-regulation, gene neighborhoods, pathway ‘cross-talk’) with one or more of the set of 14 (**Figure 1B**). The networks predicted the most highly connected, physical interactor to be ubiquitin-small subunit (40S) ribosomal protein S27A (RpS27A, Pm SeqID 173) with six interactions forming a protein-protein interaction core with another 40S ribosomal protein (RpS13, Pm SeqID 260), translation elongation (EF2, Pm SeqID 328) and initiation proteins (CG17737, Pm SeqID 72), a molecular chaperone (heat shock protein Hsp83, Pm SeqID 329), a phosphorylating enzyme (arginine kinase ArgK, Pm SeqID 200), and a protein associated with embryonic development (failed axon connections - isoform a, fax, Pm SeqID 283) (**Supplementary Table S5**). The conserved processes of protein translation, folding, and signaling appear to be central to these physical interactions (**Supplementary Table S5**). Processes represented by the physical protein-protein interactors acting outside of the RpS27A network core include protein folding (Cyp1), proton-transport (ATP binding) (ATPsynbeta), vesicle mediated transport (GTP binding) (Rab2), and carbohydrate metabolic process (Tal, GAPDH1). Examples of physical interactions included Hsp83 with one of the peptidyl-prolyl cis-trans isomerases (cyclophilin, Cyp1, Pm SeqID 189), an enzymatic protein that catalyzes protein folding, and glyceraldehyde-3-phosphate dehydrogenase (GAPDH, Pm SeqID 152) with ras-related protein Rab2 (Pm SeqID 213), a protein involved in vesicle-mediated transport and signal transduction. As expected for the ontologies and subcellular localization of the proteins represented in the physical network, there are more protein-protein functional interactions predicted among the 14 physical interactors (**Figure 1B**; **Supplementary Table S5**). Both the physical and functional protein-protein interactions among the 17 evolutionarily conserved G-interactors suggest a possible cellular coordination of processes associated with MMV G translation and protein folding and trafficking biology in insect cells.

**Figure 1.**
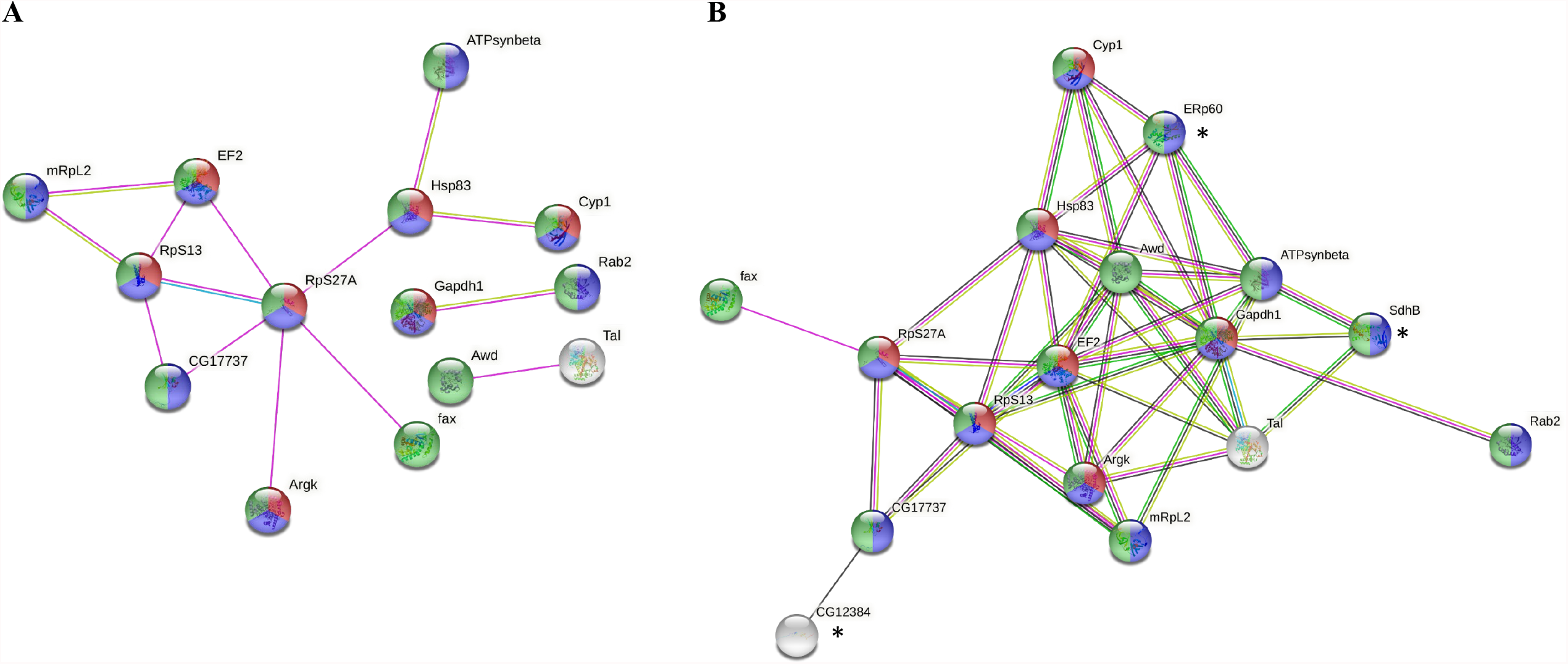
Predicted interaction networks between *Peregrinus maidis* (PM) proteins that were determined to directly interact with maize mosaic virus (MMV) glycoprotein (G) in the MbY2H assay. **A**) Physical only (direct) and **B**) physical and functional interactions between PM proteins as determined by STRING analysis (https://string-db.org/ ; Szklarczyk et al., 2021) (medium confidence = 0.4) with 100 ORF-predicted amino acid sequences (out of the 125 G-interactors) queried against a *Drosophila melanogaster* (Dmel) protein database (node IDs = Dmel gene names). Network edges (type of evidence): pink = experimental; lime green; = text-mining; blue = from curated databases; black = co-expression; green = gene neighborhood. Node colors (subcellular localization evidence): red = cytosol (part of cytoplasm, excluding organelles; blue = cytoplasm and/or organelles, excluding nucleus and plasma membrane; green = intracellular = cell components, excluding plasma membrane. Subcellular localization test (FDR): 0.0021 (cytosol), 4×10^−5^ (cytoplasm), and 0.0018 (intracellular). Refer to **Supplementary Table S5** for gene names, annotations, gene ontologies and associated PM ORF sequence IDs.

### Predicted functional sites in non-annotated MMV G-interacting proteins

Of the 57 non-annotated sequences, i.e., no blastx match or match to unknown/uncharacterized/hypothetical proteins in other species, 19 were recovered from more than one MbY2H clone (average ∼ 8 clones/sequence). Of these 19, 17 contained predicted ORFs, of which 14 had no match in public databases. ELM analysis predicted one to eight functional class sites (i.e., short linear motifs; SLiMs) per sequence (**Figure 2**), with a total of 36 represented across the 17 proteins. All 17 of these sequences (cDNA) significantly matched transcripts expressed in adult bodies or late-stage nymph guts (**Supplementary Table S4**). The SLiMs were distributed into five ELM classes indicative of function: degradation signals (ubiquitin-associated), sites of post-transcriptional modifications, docking sites, subcellular targeting sites, and ligand binding sites (**Supplementary Table S6**), and as expected, ligand binding sites were the most diverse class (24/36) (**Figure 2)**. The most enriched SLiMs across the 17 non-annotated sequences were N-degron (9), phosphotyrosine ligands bound by SH2 domains (8), PDZ domain ligands (6) and MAPK docking motifs (6), which are common motifs on proteins involved in intracellular proteolytic degradation by the ubiquitin proteasome system and signal transduction, respectively. Collectively, the putative multi-functional roles of the 17 non-annotated *P. maidis* sequences, as inferred from the SLiMs, point to nuclear transport (import and export), innate immunity, endocytosis and vesicle-mediated transport, signal transduction pathways, cell cycle, cellular communication and adhesion, protein synthesis and degradation, DNA repair and transcriptional repression (**Supplementary Table S6)**. Of potential relevance to MMV, a nuclear-replicating virus, two of these non-annotated G-interacting proteins, Pm SeqID 199 and 235, uniquely contain a predicted nuclear localization signal (NSL) and nuclear export signal (NES), respectively. These motifs align with the biology of the MMV G protein since it is known to localize to the nucleus, and the virus particle buds from the nuclear membrane (Herold and Munz, 1965; Martin and Whitfield, 2018). The NES-containing interactor (Pm SeqID 235) was the third most common protein identified in the MbY2H screen, recovered 25 times. Other SLiMs predicted from this sequence include three functional sites that indicate roles in protein trafficking and vesicle-mediated transport (Atg8 protein family ligand, adaptin binding endosome-lysosome basolateral sorting signal, and clathrin box motif). We hypothesize that this unique *P. maidis* protein may play a role in MMV-trafficking in the insect cell. The NLS containing sequence (Pm SeqID 199) was retrieved six times in the MbY2H screen and also contains predicted PDZ-ligand and phosphotyrosine-ligand motifs, both known to play roles in various cellular signaling pathways (Chi et al., 2012; Kaneko et al., 2012). Two other sequences (Pm SeqID 128 and 241) contained predicted SLiMs implicated in clathrin-mediated endocytosis (adaptin binding endosome-lysosome basolateral sorting signal and clathrin box, respectively). Ligand binding sites implicated in immunity were also predicted. The IAP (inhibitor of apoptosis)-binding motif (IBM) was found in three unique interactors, Pm SeqID 168, 203, and 285. Pm SeqID 168 was retrieved 18 times in the MbY2H screen, indicating a strong interaction with MMV G. Other ligand-binding SLiMs that indicate modulation of host response to pathogens - IRF-3, immunoreceptor tyrosine, and PDZ ligand domains - were also predicted among the set of non-annotated *P. maidis* proteins. Together with the finding that the G-interacting, non-annotated transcript sequences matched *P. maidis* sequences in previously published studies (**Supplementary Table S4**), the occurrence of putative functional sites in these sequences indicates novel insect proteins or *P. maidis*-specific proteins.

**Figure 2.**
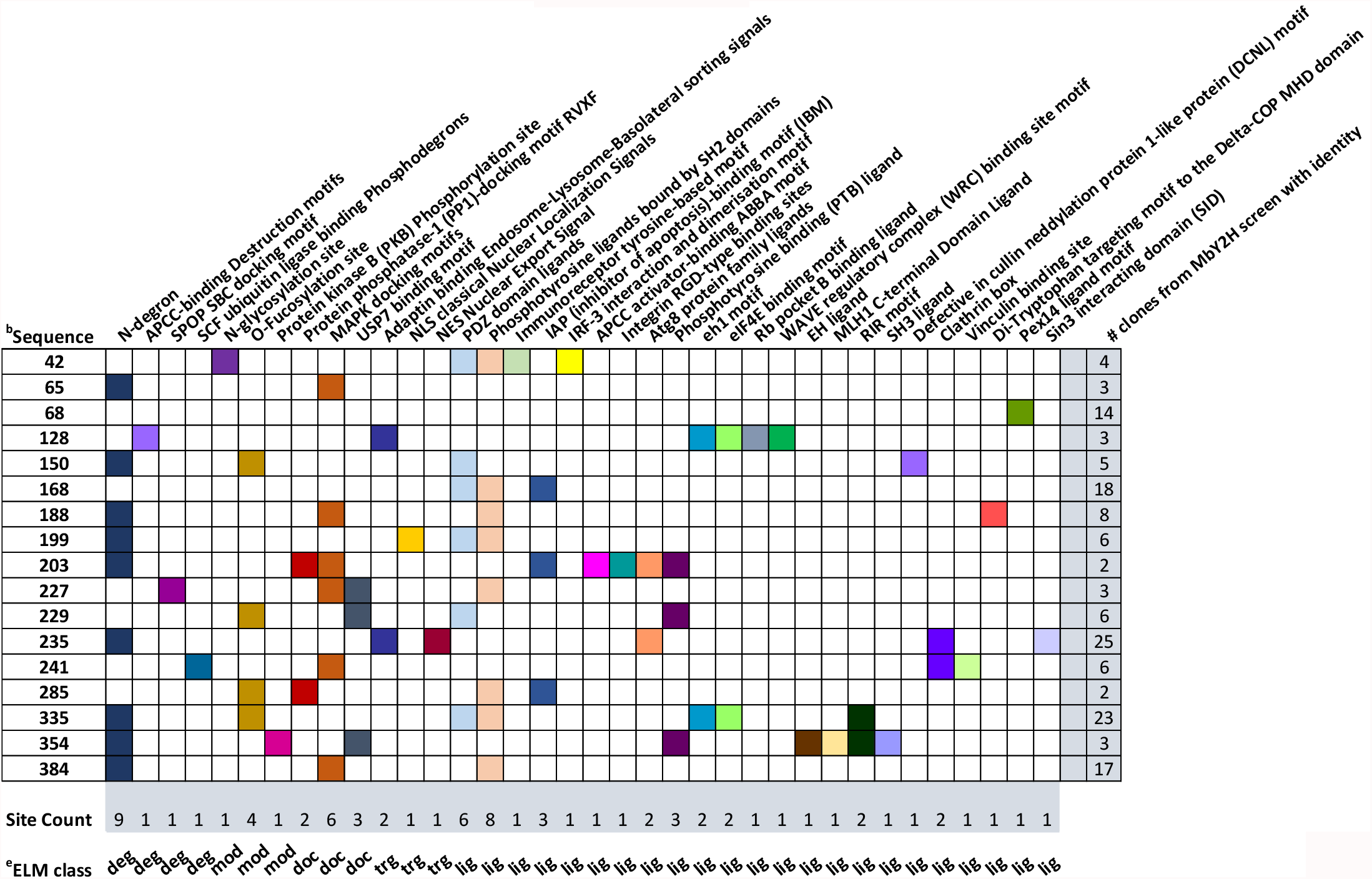
Prediction of functional sites (short linear motifs; SLiMs) in nonannotated *Peregrinus maidis* protein sequences that putatively interacted with maize mosaic virus glycoprotein (MMV-G) in the membrane bound yeast II hybrid (MbY2H) screen. Nonannotated sequences are *P. maidis* sequences with no match to publicly available sequences AND unknown, uncharacterized, or hypothetical proteins in other metazoans. SLiMs were identified using the Eukaryotic Linear Motifs (ELM) database (http://elm.eu.org/index.html; Dinkel et al., 2012); ELM analysis parameters: *Peregrinus maidis* specified organism; motif probability cut-off = 0.001 (E < 10^−4^); motifs with low conservation not reported. ^a^Sites are presented by ELM classes (types), not in order of relative position along the peptide sequence; bSubset of nonannotated sequences with at least two instances in the 432 MbY2H clone set (i.e., cluster size greater than 2); cELM class designations. Refer to **Supplementary Table S6** for functional class sites, ELM class designations, ELM ID and accessions, and functional roles of the predicted SLiMs in the dataset.

### Co-immunoprecipitation assays confirm physical interactions between MMV G and P. maidis protein candidates

In the MbY2H library screen, we identified CypA and ApoLpIII as interactors with MMV G. Cyclophilins are ubiquitous, evolutionarily well conserved proteins and CyPA functions include protein folding, trafficking, assembly, cell signaling and immunomodulation (Nigro et al., 2013). ApoLpIII is a key protein in insect hemolymph lipid transport processes and an emerging role in insect innate immunity (Weers and Ryan, 2006; Whitten et al., 2004). Bioinformatic analysis of the library suggested that these were robust interactors with G and sequences annotated as Cyp A were isolated seven times and ApoLpIII five times. To validate this result, we performed co-immunoprecipitation experiments with the target proteins, CypA and ApoLpIII, in insect cells (S2) with MMV G using an anti-FLAG affinity gel. MMV G was fused with a FLAG tag while CypA or ApoLpIII was fused with Myc tag. Before co-immunoprecipitation, successful expression of proteins in S2 cells was validated using immunoblot (data not shown). Indeed, CypA or ApoLpIII co-immunoprecipitated with MMV G as revealed in the immunoblot using a FLAG or Myc antibody (**Figure 3**).

**Figure 3.**
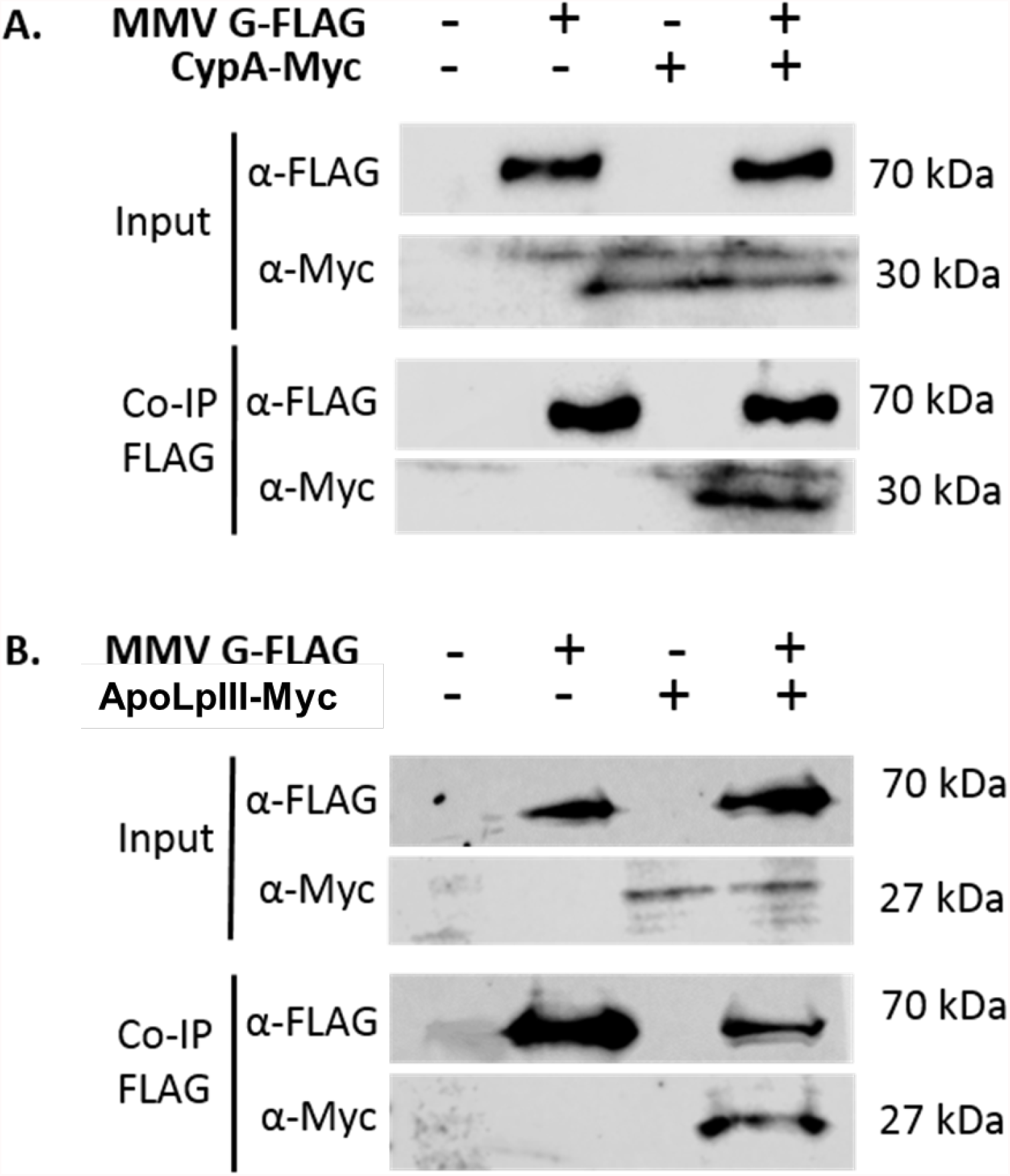
Co-immunoprecipitation of CypA or ApoLpIII and MMV G using anti-FLAG affinity gel shows that MMV G interacted with Cyclophilin A and ApoLpIII. Myc-tagged Pm proteins, CypA or ApoLpIII were co-immunoprecipitated with MMV G-FLAG using anti-FLAG affinity gel (Biotool, Huoston, TX).

### MMV G and candidate P. maidis proteins co-localize in insect cells

MMV-G-GFP expression and localization was observed in insect cell lines. In S2 cells that were transfected with MMV-G-GFP, the G fusion protein localized to the primarily to the nucleus, with some protein found in the ER and lysosomes as determined by observing similar patterns with commercially available probes for these organelles (Supplementary Figure S2). G-GFP colocalization with the lysosomal marker showed intense staining in these punctate structures. Co-expression experiments with *P. maidis* interactors and G-GFP in Sf9 cells provides insight into the location of the viral protein-insect protein interactions. The control treatment of GFP expressed alone showed protein in both the cytoplasm and the nucleus of the cell (**Figure 4**). When G-GFP was expressed alone, it matched the appearance of earlier localization patterns reported (Martin and Whitfield, 2018) with localization primarily around the nucleus with the presence of some punctate bodies around the nucleus, and the expression and localization pattern of MMV G is consistent in both S2 and Sf9 cells. When expressed in the presence of ApoLpIII and CypA, the expression of G-GFP was similar, however, ApoLpIII-RFP was also observed around the nucleus and in some cases was localized in punctate bodies near to G localization. In a few cases, it appeared the MMV-G surrounded the ApoLpIII-RFP regions. When G and CypA were co-transfected into cells, CypA was distributed throughout the cell. The punctate bodies of G-GFP in the perinuclear region could sometimes be seen in the CypA-RFP areas. In contrast, the punctate bodies of G-GFP observed in the cytoplasm did not co-localize with ApoLpIII-RFP, as ApoLpIII-RFP and MMV G were only seen in proximity near the nucleus.

**Figure 4.**
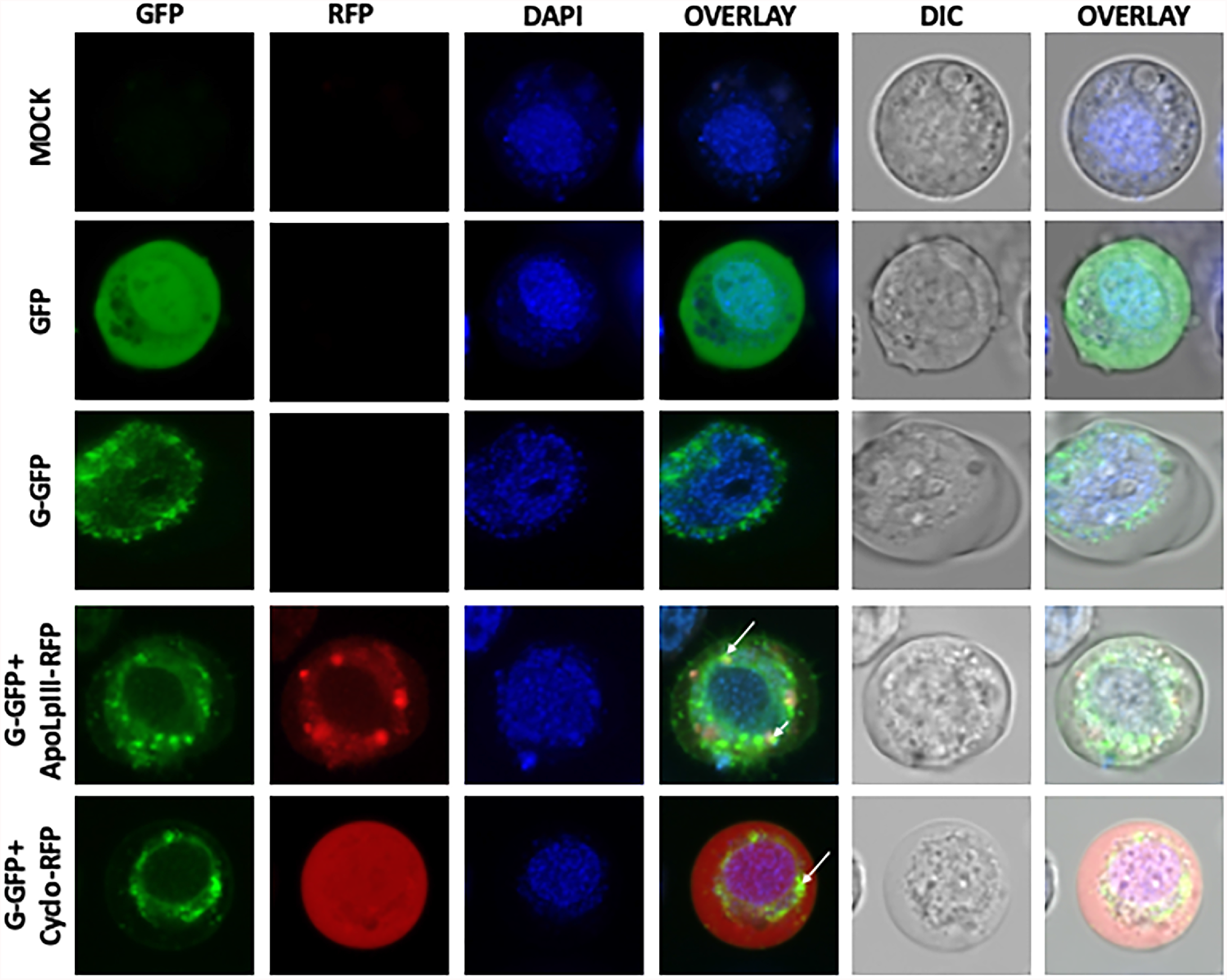
MMV G co-localizes with CypA and ApoLpIII. RFP-tagged *Peregrinus maidis* proteins were co-transfected with GFP-tagged MMV G. Two days post-transfection, the cells were visualized using laser scanning confocal microscope Zeiss LSM 880. White arrows indicate areas where the insect vector protein and G signals overlap. Scale bars are equal to 10uM.

## DISCUSSION

Rhabdovirus glycoproteins serve critical roles in host cell recognition, membrane fusion, virion packaging and movement in host cells (Albertini et al., 2012; Ammar et al., 2009; Belot et al., 2019). This study identified *P. maidis* proteins that interacted directly with MMV G protein, and provisional annotation of these interactors revealed proteins with putative roles in virus entry, membrane fusion, infection, and dissemination in the planthopper vector. Use of the split ubiquitin membrane-based yeast two hybrid system (MbY2H) enabled expression of MMV G as an integral membrane protein to capture membrane-associated interacting partners as well as cytosolic interactions. In fact, while the *P. maidis* G-interactors were predicted to localize to diverse subcellular compartments, there was an enrichment in the category of ‘integral component of a membrane.’ In line with a nuclear replicating virus, several interacting proteins contained predicted nuclear import and export motifs, including those proteins with no match to publicly available sequences (*P. maidis*-specific) or uncharacterized hypothetical proteins in other metazoans. MMV G protein is known to localize to the nucleus, and the virus particle buds from the nuclear membrane (Herold and Munz, 1965; Martin and Whitfield, 2018). Several interactors were also provisionally implicated in endo- and exocytosis and vesicle-mediated transport, a process that involves membrane fusion and intracellular, cytosolic interactions as viral genomes are shunted to the ER and Golgi for protein synthesis and post-translational modifications. In the case of the evolutionarily conserved proteins represented by a subset of the G-interactors, the physical and functional protein-protein interactions indicate a possible cellular coordination of processes associated with MMV G translation, protein folding and trafficking as well as protein degradation and turnover in insect cells.

Based on functional predictions of the *P. maidis* proteins and empirical evidence of MMV G localization (Martin and Whitfield, 2018 and this study), it appears that G is central to pathways that govern membrane fusion. The MbY2H screen revealed different proteins known for their involvement in receptor-mediated endocytosis (clathrin-mediated), a signal-dependent membrane-fusion and cargo trafficking process. Members identified in our screen included Syntaxin 18, a soluble N-ethylmaleimide sensitive factor attachment protein receptor (SNARE) protein, small GTPase Rab2, nucleoside diphosphate kinases (NDK), and E3 ubiquitin protein ligase MARCH3 (Membrane-associated RING-CH). STX18 is known to function in trafficking vesicles within the cell, into or out of the ER (Hatsuzawa et al., 2000; Teng et al 2001). This SNARE protein was documented to interact with bovine papillomavirus type 1 L2 capsid protein (Bossis et al., 2005) and it is hypothesized to be involved in ER-mediated phagocytosis by regulating specific and direct fusion of the ER and plasma or phagosomal membranes (Hatsuzawa et al., 2000). NDK - which serves a role in normal presynaptic function (Krishnan et al., 2001) - may cooperate with STX18 to play essential roles in the neurotropic infection in *P. maidis*. Indeed, seminal work documented extensive MMV infection in the nervous system tissues of *P. maidis* (E. D. Ammar 2008). Similar to STX 18, small GTPase Rab2 plays a critical role in membrane trafficking events and is required for membrane transport in the early secretory pathway (Tisdale et al., 1996). Dissecting the roles of STX18, Rab2 and NDK in the planthopper vector warrants future exploration to distinguish their possible roles in host cell entry/exit of MMV and systemic infection in the insect host.

Nearly half of the putative G-interacting, *P. maidis* proteins appear to be novel (specific to *P. maidis*) or serve unknown functions in other animals. It is not uncommon for non-model insect transcriptomes to contain numerous non-annotated transcripts with predicted protein features (Martin et al., 2017; Thorpe et al., 2016; Schneweis et al., 2017; Xeu et al., 2010). In this study, a cross-comparison with published *P. maidis* transcriptomes derived from RNAseq (Martin et al., 2017) and Sanger EST sequencing (Whitfield et al., 2011) provided confidence in the occurrence of these apparent *P. maidis*-specific, G-interactors. Moreover, non-annotated sequences recovered from more than one MbY2H clone contained myriad putative functional sites indicative of particular cellular roles, many of which are consistent with the provisional functional roles of the known proteins identified by the screen, e.g. endocytosis, vesicle-mediated transport, protein degradation and turnover, and signal transduction. Of particular interest, the second most common ligand binding motif in our analysis was the PDZ-domain ligand binding site. PDZ domains are well-known protein-protein interaction modules which can be a target by the viral PDZ-binding motif present in MMV G (Martin et al., 2017). Future mutational analysis and/or genome-editing of PDZ-domain and other ligand-binding motifs in the set of non-annotated, G-interacting proteins may reveal novel vector proteins associated with MMV transmission biology.

Moving beyond *in silico* sequence analysis, two *P. maidis* G-interactors were advanced forward for their reported role in innate immunity and hemolymph activities (apolipophorin III; ApoLpIII) and implication in other vector-plant virus systems (peptidyl-prolyl cis-trans isomerase; Cyclophilin A (CypA). In this study, ApoLpIII and CypA sequences were recovered multiple times from the MbY2H screen and the direct interaction was robustly confirmed by co-immunoprecipitation assays. Furthermore, ApoLpIII and CypA co-localized with MMV G in a heterologous insect cell system, with differing patterns of distribution in relation to the viral glycoprotein.

ApoLpIII is an insect apolipoprotein with documented roles in lipid transport, pathogen pattern recognition, and multicellular encapsulation reactions as a component of the innate immune response (Niere et al., 1999; 2001; Weers et al., 2003). ApoLpIII lipid transport proteins are abundant in the hemolymph, and found in hemocytes, eggs, and the fat body of diverse insects (Weers et al., 2006). MMV has been detected in planthopper hemocytes and may disseminate to other tissues following a hemocyte route (Ammar and Hogenhout 2008; Yao et al., 2019), thus it is possible that ApoLpIII in the *P. maidis* hemolymph could be exploited for virus particle transport. In a differential hybridization screen of a cDNA library generated from condemned (destined to die) intersegmental muscles of *Manduca sexta*, ApoLpIII was found to be upregulated, suggesting its role in programmed cell death (Sun et al., 1995). In contrast, *Anopheles stephensi* challenged with *Plasmodium berghei* showed an increase in ApoLpIII expression in midguts and silencing of this gene resulted in a decrease in oocycst numbers suggesting that ApoLpIII is a positive regulator of pathogen development (Dhawan et al., 2017). Determination of whether the direct interaction between MMV G and ApoLpIII initiates a pro- or antiviral response in the insect vector will be the focus of functional assays.

Cyclophilins, also known as peptidyl-prolyl cis-trans-isomerases, are ubiquitous proteins involved in multiple biological processes, including protein folding and trafficking, cell signaling, immune responses and numerous interactions between plant viruses in their insect and plant hosts (Kanakala and Ghanim, 2016; Kovalev and Nagy, 2013; Kumari et al., 2013; Tamborindeguy et al., 2013). While the functional role of *P. maidis* CypA in MMV has not been determined, the direct interaction between CypA and viruses appears to be conserved. For the negative-strand RNA virus tomato spotted wilt virus (TSWV), the thrips vector cyclophilin was shown to bind to the virus attachment protein, GN, and the interaction occurred in yeast and insect cells (Badillo Vargas et al., 2019). For viruses transmitted by whiteflies and aphids, various cyclophilins have also been identified to interact with purified virus or capsid proteins (Kanakala and Ghanim, 2016; Tamborindeguy et al., 2013). For some other systems, the functional role of cyclophilin A has been determined. CypA is essential for trafficking viral (HIV-1) capsid structures to the perinuclear region and inhibition of the binding of the capsid to cypA decreased infectivity of the virus (Sokolskaja et al., 2004; Zhong et al., 2021). The CypA binding to capsid in the cytoplasm prevents premature uncoating of the nucleocapsid and enables the complex to reach the perinuclear region. In insect cells, we observed a widespread distribution of *P. maidis* CypA and some regions of overlap in localization with G. As a nuclear-replicating virus, MMV has a similar requirement for transport to the nucleus for uncoating and replication. For another member of the Rhabdoviridae, VSV, Bose et al. (2003) identified CypA as one of the cellular co-factors critically required for VSV-NJ replication but not for replication of the serologically distinct VSV-IND. Another role for cyclophilin comes from the pathogen, *Trypanosoma cruzi*, and the reduviid vector. In this interaction, the parasite secretes cyclophilin in the intestine of the vector where it serves to protect the parasite from insect-host secreted antimicrobial peptides (Kulkarni et al., 2013). Intriguingly, we also identified an antimicrobial peptide, lugensin, as an interactor with MMV G. It may be that there is a coordinated response between glycoprotein binding proteins during virus infection of the insect vector. If cyclophilin serves a protective role in the insect gut, down-regulation of expression may make the virus a better target for antimicrobial responses like lugensin.

## CONCLUDING REMARKS

The identification of host proteins interacting with G provided new knowledge about the rhabdovirus-vector interactome. Physical interaction of MMV G with proteins implicated in membrane fusion can be used to better understand the events leading to successful acquisition, dissemination and replication of MMV in the insect-vector host. Because MMV G can interact with numerous vector proteins with diverse functions, it is possible that a ubiquitous molecule or more than one molecule can serve as the virus receptor depending on the tissue being invaded. The identification of proteins implicated in the host’s synaptic physiology can be further investigated to better understand how this plant virus is capable of invading and infecting the nervous system yet does not cause obvious pathogenic effects. The verified interaction of G with CypA and ApoLIII may suggest a means towards developing RNAi-based biotechnology products for reducing virus transmission and/or controlling the highly destructive corn planthopper.

## Supporting information

Supplementary Fasta Sequence File

## AUTHOR CONTRIBUTIONS

Conceived and designed the experiments: AEW, KA. Performed the experiments: KA, KM. Bioinformatic analysis: DR, KA. Analyzed the data: KA, DR, AEW. Wrote the paper: KA, DR, AEW.

## ACKNOWLEDGMENTS

We thank members of the Plant-Virus-Vector Interactions Lab team for thoughtful discussions throughout the course of the project.

## SUPPLEMENTARY MATERIALS

**Supplementary Fasta Sequence File:** 125 non-redundant *P. maidis* sequences from MbY2H protein-protein interaction screen with MMV-G (.txt)

**Supplementary Table S1:** Primer sequences designed for cloning and sequencing (.docx)

**Supplementary Table S2:** MbY2H host strain genotypes (.docx)

**Supplementary Table S3:** Annotations and gene ontologies for the 125 non-redundant *P. maidis* sequences (.xlxs)

**Supplementary Table S4:** Significant BLASTn matches between the 125 non-redundant *P. maidis* sequences (cDNA) and transcripts expressed in the gut (nymph) and whole bodies (adult) *P. maidis* (.xlxs)

**Supplementary Table S5:** Annotations of conserved proteins mapped to *Drosophila melanogaster* proteome sequences as part of the STRING protein-protein interaction network analysis for the 125 non-redundant *P. maidis* sequence dataset (.xlxs)

**Supplementary Table S6:** ELM functional sites, classes, accessions and annotations for non-annotated putative proteins in the 125 non-redundant *P. maidis* sequence dataset (.xlxs)

**Supplementary Figure S1:** Pie charts depicting distribution of the 125 non-redundant *P. maidis* sequences into gene ontologies

**Supplementary Figure S2:** Immunolocalization of MMV G protein in insect cells (*D. melanogaster* S2 cell line)

