## Supplementary Table S1 for "Identification of interacting proteins of maize mosaic virus glycoprotein in its vector, *Peregrinus maidis*"

**Supplementary Table S1**. Primers designed and used for cloning and sequencing in the study.

| **Primer sequence name** | **Primer use** | **Sequence (5’ – 3’)** |
| --- | --- | --- |
| MMV G | Directional cloning into pBT3-SUC | F: ATTAACAAGGCCATTACGGCCATGTCTCTCATTCACCT  CATCTACTTC  R: AACTGATTGGCCGAGGCGGCCCCTGACCTTACCCATTC  TATATTTGT |
| SUC-MMVG | Bait construction | F:ATTAACAAGGCCATTACGGCCACACCAGCATTGACTTATGGCAACCTG  R:AACTGATTGGCCGAGGCGGCCCCTGACCTTACCCATTCTATATTTGTGGG |
| pBT3-SUC | Sequencing of the bait construct | F: TGGCATGCATGTGCTCTG  R: GTAAGGTGGACTCCTTCT |
| pPR3-N | Sequencing of preys | F: GTCGAAAATTCAAGACAAGG  R: AAGCGTGACATAACTAATTAC |
| HSP | Sequencing of clones for S2 co-localization study | F:TATAAATAGAGGCGCTTCGT  R: CTTCGGGCATGGCGGACTTG |
| GFP |  |  |
| attb1 |  | F: ACAAGTTTGTACAAAAAAGCAGGCT  R: ACCCAGCTTTCTTGTACAAAGTGGT |
| attb2 |  |  |
| MMV G | pENTR cloning for Gateway recombination into insect expression vector (*Drosophila* S2 cells) | F: CACCATGTCTCTCATTCACCCTCATCTAC  R:TGACCTTACCCATTCTATATTTTG |
| Apolipophorin III |  | F:CACCATGGCAAACTATTGTACTGTTTAC  R: GTGTTGATGACCTTCATGACCTTC |
| Cyclophilin A |  | F: CACCATGGCACGTTCAAAGGTATAC  R: GAGCTGACCGCAGTCGGCGATTG |
