## Supplementary Table S2 for "Identification of interacting proteins of maize mosaic virus glycoprotein in its vector, *Peregrinus maidis*"

**Supplementary Table S2.** Yeast and bacterial host strains used in the study.

| **Host strain** | **Genotype** |
| --- | --- |
| NMY51 *(Saccharomyces cerevisiae)* | MATahis3Δ200 trp1-901 leu2-3,112 ade2 LYS2::(lexAop)4-HIS3 ura3::(lexAop)8-lacZ ade2::(lexAop)8-ADE2 GAL4 |
| XL1-Blue *(Escherichia coli)* | recA1 endA1 gyrA96 thi-1 hsdR17 supE44 relA1 lac [F´ proAB lacIq Z∆M15 Tn10 (Tetr) |
| DH5α (*Escherichia coli)* | dlacZ Delta M15 Delta(lacZYA-argF) U169 recA1 endA1 hsdR17(rK-mK+) supE44 thi-1 gyrA96 relA1 |
