## Supplementary Figure S1 for "Identification of interacting proteins of maize mosaic virus glycoprotein in its vector, *Peregrinus maidis*"

**Supplementary Figure 1. Distribution of MMV-G-interacting *Peregrinus maidis* nonredundant sequences into gene ontologies (GO) by A) biological process, B) molecular function, and C) cellular localization.** Multilevel GO charts depict sequences in the most terminal node (most specific GO) of the hierarchical GO graph.

A.

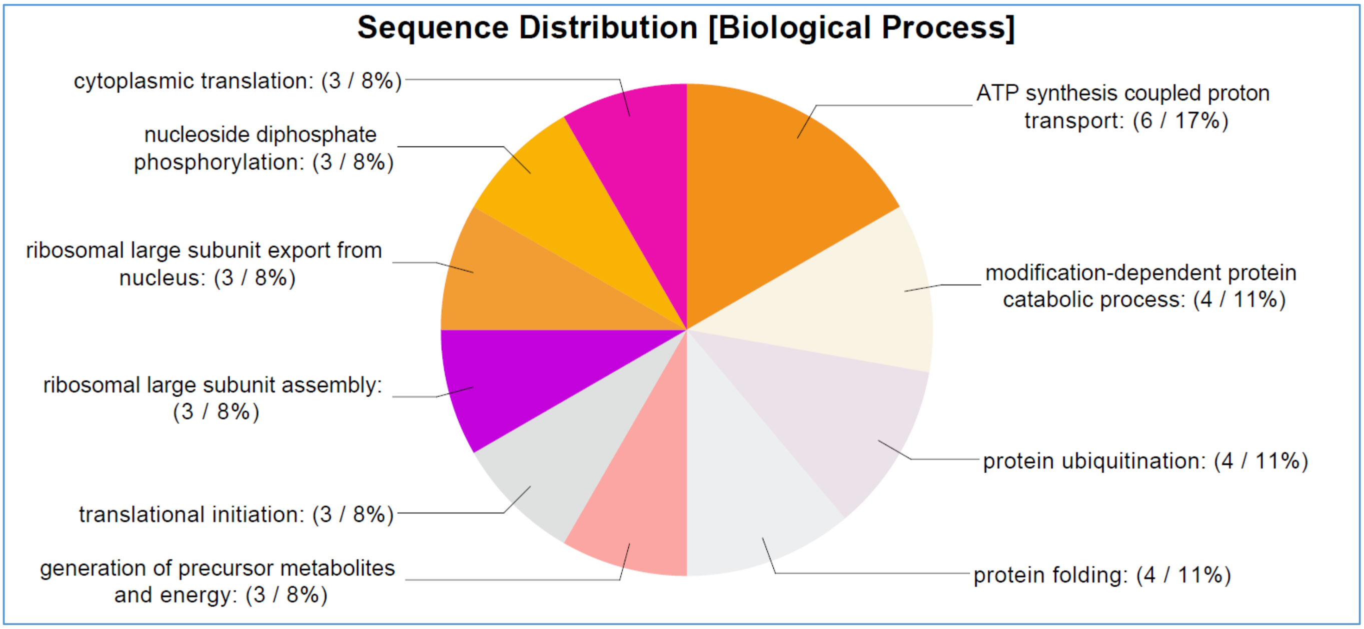


B.

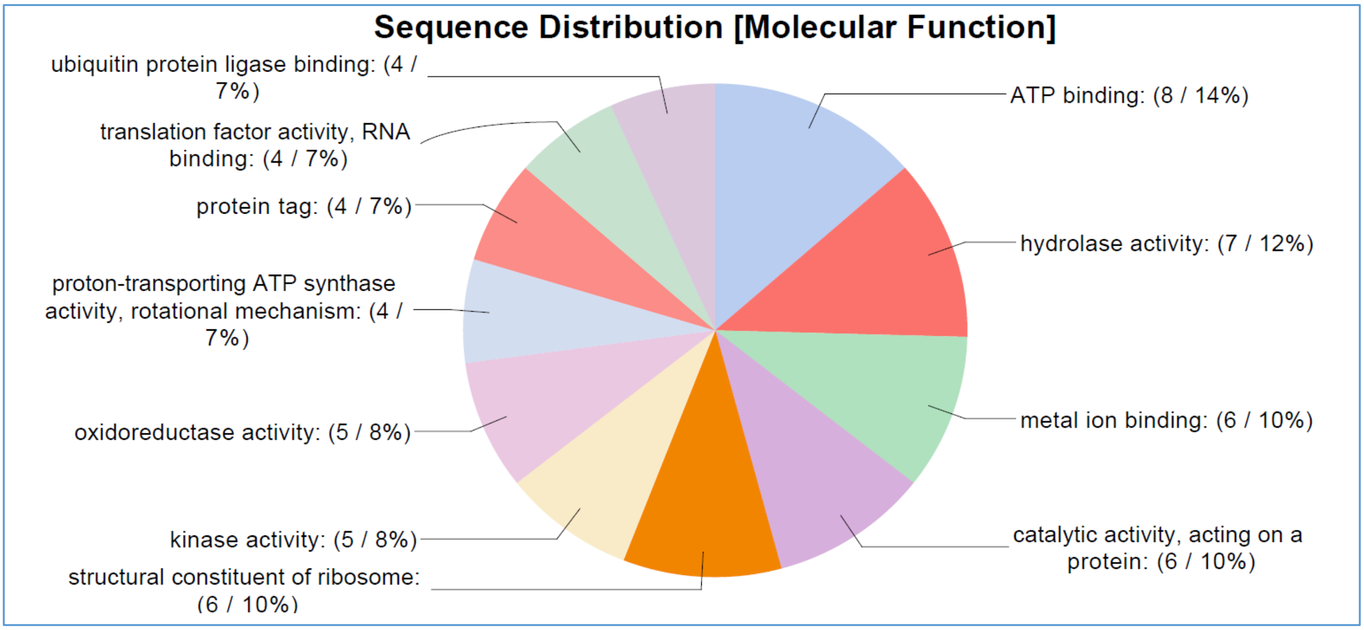


C.

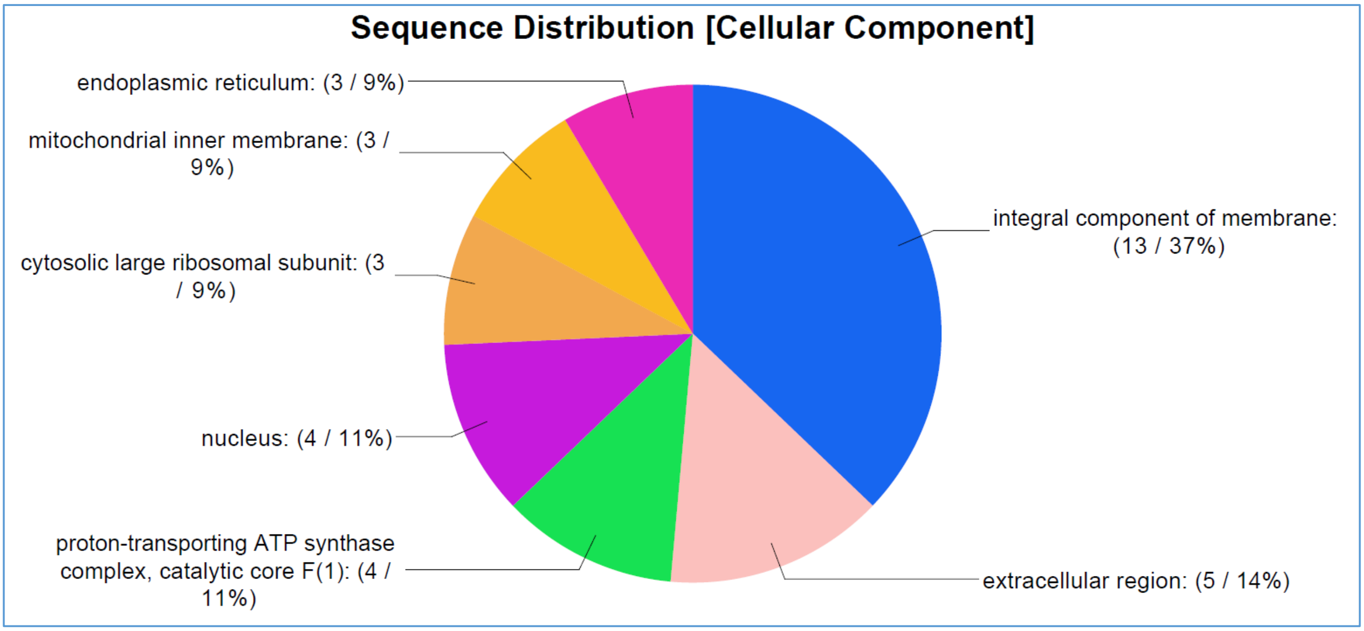
