## Supplementary Figure S2 for "Identification of interacting proteins of maize mosaic virus glycoprotein in its vector, *Peregrinus maidis*"

**Supplementary Figure S2.** MMV G localized in the ER and lysosomes. S2 cells were transfected with MMV-G-GFP and 2 days post transfection (2 dpt), the cells were visualized using laser scanning confocal microscope Zeiss LSM 780. Cells were visualized in multiple channels corresponding to emission/excitation wavelengths for eGFP and dyes. Organelle dyes, LysoTracker Red DND-99 is a red-fluorescent dye used to stain lysosomes and ERTracker Blue-White DPX is selective for the endoplasmic reticulum (ER). Multiple channels corresponding to emission/excitation wavelengths for the auto fluorescent tags and organelle markers were used to locate MMV G, lysosomes and ER in the cell respectively. MMV G appears to localize in lysosomes and the ER. Confocal images shown have 2um scale.

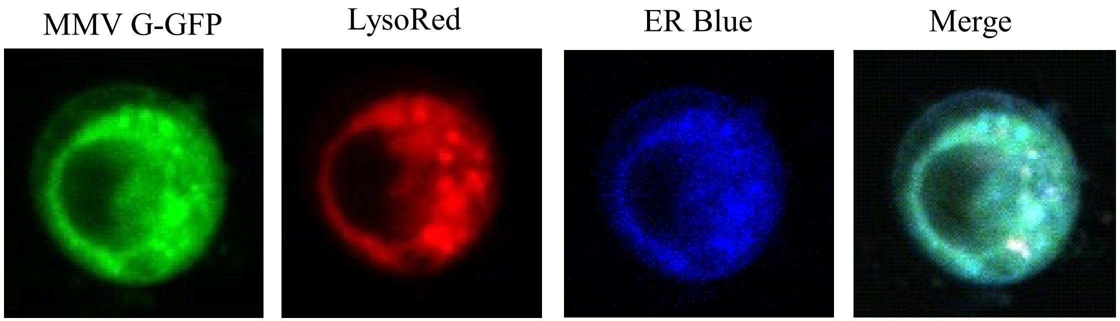
